# The emerging medium-scale: Prioritising fisheries management for elasmobranch conservation

**DOI:** 10.64898/2026.09.02.748653

**Authors:** Mayuri Chopra, Divya Karnad, Gwilym Rowlands, Guy M. W. Stevens, Mohanraj T., Katrina J. Davis

**Affiliations:** University of Oxford, Life and Mind building, South Parks Road, Oxford OX1 3SZ, UK; The Manta Trust, Catemwood House, Norwood Lane, Corscombe, Dorset, DT2 0NT, UK; Centre for Ecology and Conservation, University of Exeter, Penryn Campus, TR10 9FE, UK; Department of Environmental Studies, Ashoka University, Plot no 2, Rajiv Gandhi Education City, Rai, Sonipat, Haryana 131029, India; Ocean and Earth Science, University of Southampton, National Oceanographic Centre Southampton, Southampton, SO14 3ZH, UK; Marine Eco-biological Research Centre, Tuticorin 628001, India

**Keywords:** catch per unit effort, fisheries management, mobula, small-scale fisheries, manta ray, devil ray

## Abstract

Fishing pressure has substantially increased over time and elasmobranchs are among the most vulnerable marine taxa impacted. ‘Small-scale’ and ‘large-scale’ are terms frequently used to describe fisheries, yet definitions used across the globe are inconsistent. The global impact of small-scale fisheries on elasmobranch decline is increasingly evident. However, ambiguous vessel-capacity classifications undermine effective conservation management. Technological advancements have led to the emergence of medium-capacity vessels that sit between small and large-scale classifications, creating regulatory gaps that increase bycatch impacts on vulnerable elasmobranchs. Beyond vessel size, the gear specificities and practices also affect interaction with species caught incidentally, and can alter bycatch risk for vulnerable elasmobranchs. Manta and devil rays (collectively, mobulids) represent a pelagic ray group that includes species listed as Critically Endangered with high risk of extinction, largely due to bycatch threats. To prioritise fisheries management for conservation, we assess the contribution of fishing gear specifications and practices to mobulid bycatch as a case study in their largest fishery globally. We use a Huber-robust extension to a Hurdle negative binomial model to assess the impact of fishery characteristics and operational specificities on mobulid bycatch risk. We find that within the Food and Agriculture Organization (FAO) of the United Nations’ broad international fleet classification of small-scale vessels based on overall length (≤24 m), the ‘medium-capacity vessels’ in India are twice as likely to catch mobulids than small-scale vessels. Further, increased fishing intensity, measured in five-day increments, doubles mobulid bycatch risk. Our results indicate the emergence of a distinct medium-capacity fishery class in rapidly developing blue-economy nations, shaped by rising technological capacity and exemptions from large-scale management measures. We propose a comprehensive re-conceptualisation of small-scale fisheries to ensure that advanced medium-capacity vessels are subject to management measures commensurate with their bycatch impact. Our recommended re-conceptualisation is complementary to FAO’s SSF characterisation matrix and would promote fishery-level sustainability and enable bycatch risk mitigation for elasmobranchs, including manta and devil rays. Further, for congruence between elasmobranch demography and regulations, we recommend that fisheries management measures distinguish vessel capacities for elasmobranch bycatch risk and apply biologically relevant regulations to the associated fisheries scale.

## 3. Introduction

Overfishing puts disproportionate pressure on vulnerable marine taxa, particularly elasmobranchs (sharks, rays, and skates), constrained by their capacity for population recovery (Stevens et al., 2000; Frisk et al., 2005; Simpfendorfer and Kyne, 2009). As fishing intensity has accelerated over the past decade, more than one-third of all elasmobranch species now exhibit significant population declines and elevated extinction risk (Davidson et al., 2016; Jabado et al. 2018; Dulvy et al., 2021). Elasmobranch stocks are especially vulnerable to overfishing due to their slow life histories characterised by low fecundities and growth rates (Frisk et al., 2005; Dulvy et al., 2014a; Quetglas et al., 2016; Chopra et al. 2026b). Over 90 of the 1,213 known living species of elasmobranchs (Jabado et al., 2024) are currently classified as Critically Endangered on the International Union for Conservation of Nature (IUCN) Red List of Threatened Species (IUCN, 2025), representing one of the most acute biodiversity crises in global marine systems. Weak fisheries regulatory frameworks, combined with expanding fleet sizes, exacerbate the overexploitation of vulnerable elasmobranchs already on the brink of extinction (Pacoureau et al., 2021).

A major cause of ineffective fisheries regulatory frameworks is fishery scale classification discrepancies (Laxe, 2010; Bartlett et al., 2025; Smith and Basurto, 2019). Large-scale fishery (LSF) and small-scale fishery (SSF) definitions are fluid globally and vary across developing and developed nations, as well as among international organisations (Thermes et al., 2023; Samad et al., 2025). The Food and Agricultural Organization of the United Nations (FAO) does not recognise a universal definition of small-scale fisheries but broadly classifies vessels using vessel overall length, gross tonnage, and engine power; however, the relevance of these metrics varies with fishing gear. For example, engine power is highly relevant for trawlers but not for gillnetters (Thermes et al., 2023). The overall vessel length over 24 m is context-dependent and not consistently applicable to fishing fleets in the Global North and the South (FAO, 2020; Thermes et al., 2023; Smith and Basurto, 2019). More importantly, the classification of fishery scale discrepancies is not limited to vessel size regulations, but extends to ineffectiveness in associated policy measures, such as mesh size regulation, minimum landing size for species, spatial bans, and seasonal closures (Davies et al., 2018). This is problematic because these policy measures that regulate fisheries also impact bycatch risk for vulnerable species. By extension, fishery capacity classification discrepancies exacerbate management complexities for vulnerable elasmobranchs, threatening their conservation (Temple et al., 2018; Smith and Basurto, 2019).

LSFs have historically been a major driver of overfishing and elasmobranch decline (Pacoureau et al., 2021). Large-scale vessels often operate in the high seas or the Area Beyond National Jurisdiction (ABNJ), where policy implementation and catch regulation are limited (Neat, 2025). Despite numerous resolutions for bycatch mitigation, over 75% of policies adopted by Regional Fishery Management Organisations (RFMOs) are unlikely to avoid or minimise elasmobranch bycatch (Cronin et al., 2023). By contrast, SSFs, which comprise 40% of global fisheries catches and support 2.3 billion people, were historically assumed to have a lower impact on elasmobranch populations (Basurto et al., 2025). However, SSFs’ increasing fishing pressure and large cumulative impact are now recognised globally (Fernando and Stewart, 2021; Di Lorenzo et al., 2022), and the unregulated expansion of SSFs has been identified as a concern for both long-term fishery sustainability and the conservation of large, slow-growing marine vertebrates (Alfaro-Shigueto et al., 2010). As a result, differences in vessel capacity and operational range across fishery scales can translate into greater interactions between vessels and vulnerable elasmobranch populations (Lyons et al., 2013). This risk is particularly of concern within SSFs that operate predominantly in data-poor coastal regions where bycatch impact assessments remain challenging (Temple et al., 2019; Pita et al., 2019).

Within SSFs, advanced mechanised vessels are categorised as small-scale despite their enhanced capacities from modern gear materials and technological advancements such as electronic gill net retrievers, engine modifications, and fuel-efficient infrastructure (Kaur and Datta, 2021). These advanced mechanised vessels gain an advantage from their small overall length (OAL), which exempts them from certain LSF management measures such as spatiotemporal bans (Gunakar et al., 2017). Additionally, they benefit from the subsidies intended for subsistence fisheries (Sumaila et al., 2019). Their comparatively higher fuel efficiency relative to large-scale vessels reduces operating costs, which further strengthens this advantage (Asokan et al., 2023). Together, these regulatory, financial, and operational advantages allow such vessels to function beyond the ‘small-scale’ their classification implies. As a result, ambiguous capacity classifications mean that government policies designed to support subsistence fisheries may inadvertently subsidise medium-scale fleet fishing effort at levels that are not optimal from both welfare and conservation perspectives. Therefore, maintaining their classification as small-scale creates regulatory blind spots, allowing technologically advanced vessels to remain governed under SSF frameworks. This misalignment has negative implications for elasmobranch sustainability, particularly in nations advancing blue economy agendas (Jadhav, 2018).

Beyond vessel size, fishing practices and gear characteristics further mediate how different fleets interact with non-target species. Fishing intensities that enable varied spatial zones of fishing can impact the species composition and the abundance of threatened bycatch species (Lewison et al., 2009; Stewart et al., 2010; Fauconnet et al., 2023). Additionally, fishing gear specifications, such as mesh size, actual fishing height (based on buoyancy devices used), and net hanging ratio (a measure of the maximum net stretch), can modify the tangling effect of the net on vulnerable bycatch species (Northridge et al., 2017). Gear soak time and depth of fishing have also been correlated with an increased bycatch risk for elasmobranchs (Morgan et al., 2010; Lyons et al., 2013). Therefore, to achieve stronger fisheries management regulation, the identification of fishery practices and fleet demographics that heavily contribute to bycatch is necessary to streamline conservation policy and thereby increase the efficiency of conservation measures.

In this case study, we identify the contribution of fishery characteristics to manta and devil ray catch in India, which hosts their largest fishery globally (Laglbauer et al., 2026), to demonstrate the high-capacity fleet contribution to elasmobranch bycatch. India provides a particularly relevant case study, as its fisheries are dominated by small-scale fleets undergoing rapid mechanisation and emerging as an advanced fishing fleet (Jadhav, 2018; Muruganandam et al., 2019). We take manta and devil rays as our case study species due to their populations facing up to 99% declines globally (Dulvy et al., 2014b; Pardo et al. 2016, Barrowclift et al., 2025; Jabado et al., 2025a, 2025b, 2025c; CITES, 2025). International policies, including CITES, are dependent upon effective national fisheries management. This management is globally low, as exemplified by extensive overfishing of mobulids (Palacios et al., 2025; Laglbauer et al., 2026). In this context, we test the following hypotheses to evaluate how fleet capacity and fishing practices interact to shape mobulid bycatch risk: (H1) Medium-scale vessels have the highest mobulid bycatch, (H2) Large-scale vessels have the highest mobulid bycatch per unit effort, (H3) Mobulid mean catch per unit effort increases with larger mesh sizes, increasing distance from the coast, and greater depth of fishing, (H4) The encounter probability of mobulid bycatch per unit effort is higher for medium-scale than for small-scale vessels, and (H5) The abundance probability of mobulid bycatch per unit effort is higher for advanced vessel sizes and fishery practices. This study is the first to identify the fishing fleet segment most important for mobulid conservation action in India, and is the most detailed catch per unit effort analysis of mobulids in the case study area to date. Our results highlight that capacity differences within small and large-scale fisheries and associated practices affect vulnerable elasmobranchs differently. For global applicability, we also highlight the significant differences in bycatch contribution between vessel types and fishery practices, which we propose are leading to the emergence of a separate capacity class of ‘medium-scale fisheries’ that is distinct from SSFs in nations rapidly developing their blue-economies.

## 4. Methods

### 4.1 Study site

We focus on India as a case study due to its intensive marine fisheries and importance in global mobulid conservation (O’Malley et al., 2017; Palacios et al., 2025), with severe reported declines in manta and devil rays alongside a multi-fold increase in fisheries pressure (Chopra et al., 2026a). India’s blue economy has seen a rapid increase, with a large proportion of relatively advanced mechanised vessels still categorised as ‘small-scale’. The reported catch in Indian fisheries has grown six times since the nation’s independence in 1947 (Ansell, 2020). It is the fourth largest global economy and among the fastest growing worldwide (Lee and Song, 2025), with blue economy forming a significant component of its economic development strategy (Karuppiah et al., 2025). The impact of middle-scale vessel emergence is likely reflected in global fishery contributions, as India is the third largest capture fisheries producer (FAO, 2024). We identified fisheries management priorities for mobulid conservation in the southeastern Indian state of Tamil Nadu (Figure 1) due to its reportedly high landings (Mohanraj et al., 2024; Chopra et al., 2026a).

**Figure 1.**
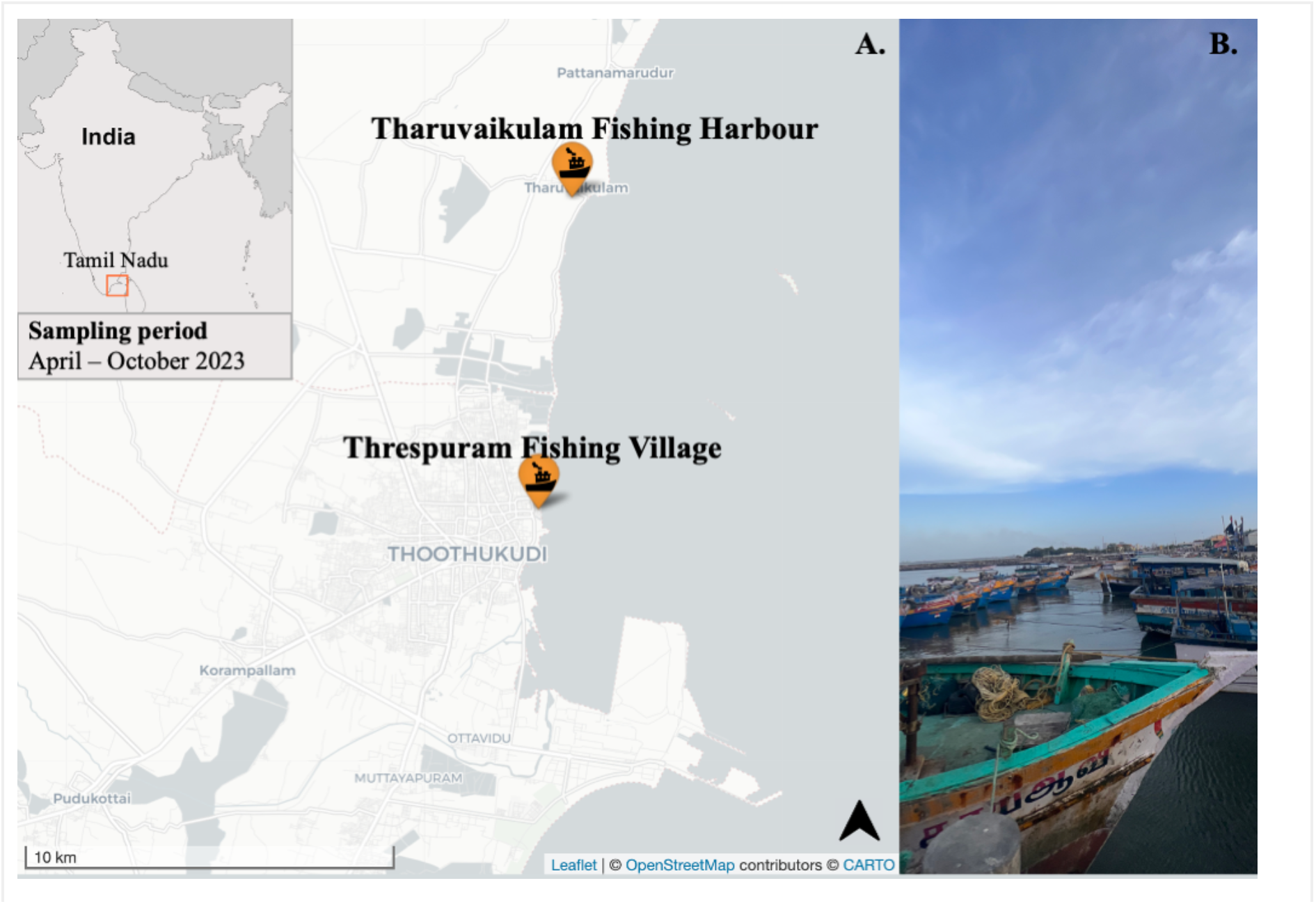
Study area to identify fishery management priorities for mobulid bycatch mitigation. The study area is in Thoothukudi District with A. Sampling locations at Threspuram Fishing Village and Tharuvaikulam Fishing Harbour indicated, and B. Landscape view of a sampling location jetty in Thoothukudi District, Tamil Nadu, India

To identify fisheries management priorities, we assessed the contribution of fishery characteristics to mobulid landings using data collected by Chopra et al. (2026a). As reported in Chopra et al., (2026a), we collected data on manta and devil ray (hereafter, mobulids) landings in 2023 at two fish landing sites in the Thoothukudi District: Threspuram Fishing Village (8.8165° N, 78.1625° E) and Tharuvaikulam Fishing Harbour (8.8922° N, 78.1707° E) (Figure 1). We focused on Tamil Nadu (Figure 1) because it contributes 86% of all east coast mobulid landings in India (Kizhakudan et al., 2015; Nair et al. 2015). Within Tamil Nadu, we focused on Thoothukudi, the state’s third-highest fishery landings district (CMFRI, 2024), which also reports high mobulid landings (Couturier et al., 2012; Sivadas et al., 2013).

### 4.2 Data collection

As described by Chopra et al. (2026a), the landings data were collected from April to October in 2023, prioritising Thoothukudi’s peak tuna fishing season (June to September), when mobulid landings occur in gill net fisheries targeting tuna and other pelagic fish (Kumar, 2017). To assess the contribution of fishery characteristics to mobulid landings, we collected data on boat size (ft), mesh size (mm), distance of fishing from the coast (nautical miles), depth of fishing gear (m), net soak time (hrs), and days spent fishing on a trip.

### 4.3 Data analysis

We collected the fishery characteristic variables in ordered categories, coded as numeric. To ensure uniformity in data type, we transformed soak hours from continuous numeric values into ordered categories, and boat size categories were classified as small, medium, and large-scale based on their overall length in metres (OAL) (Thermes et al., 2023). These classifications were supported by the national policies wherein: the Indian Marine Policy (India G.O., 2004) defines marine fish resources classified into subsistence fishing (<12 m), small-scale fishing (12-20 m) and large-scale fishing (>20 m); whereas the Indian Marine Fisheries Act (India G.O., 2021) categorises vessels as <15 m or >15 m overall length for penalties (applied to both mechanised and motorised vessels). Therefore, all vessels >21 m were classified as large-scale; and those under 15 m were classified small-scale. The remaining size class in between the small and large-scale classification was categorised as medium-scale (15-21 m). To ensure consistent handling of cases with respondent variation, e.g., different boat sizes reported for the same vessel (n = 10; 5.9% of vessels), we applied a uniform set of decision rules across all affected vessels (Supplementary Information 1). We analysed all data using RStudio version 2024.12.0+467 (R Core team, 2024). We used packages dplyr (Yarberry, 2021), lubridate (Mailund, 2019), tidyr (Wickham and Wickham, 2017), pscl (Zeileis et al., 2008), and car (Fox et al., 2012) for data processing and analysis, and factoextra (Kassambara and Mundt, 2020) and ggplot2 (Wickham et al., 2016) for data visualisation

#### 4.3.1 Mobulid total catch and catch per unit effort

To estimate the contribution of fishery characteristics to total mobulid landings and to test hypotheses (H1) – (H5) (Table 1), we aggregated total observed mobulid catch in each fishery characteristic category. To estimate mobulid catch per unit effort (CPUE), we treated a single fishing trip as the standard measure of effort, consistent with previous literature (Davie et al., 2015; Leitão et al., 2022). As our data only describes the positive occurrences of mobulid bycatch, we needed data imputation for the zero occurrences of mobulid bycatch. To assess the impact of fishery characteristics on mobulid catch per unit effort, we imputed the unobserved trips (with zero bycatch of mobulids) for identified fishing vessels in our dataset. Exercising caution, we did not impute any trips for the vessel fleet that were never observed catching mobulids in our sampling period, as we had no information on them and a proportion of these vessels could also be inactive. To impute the unobserved zero mobulid bycatch trips of identified vessels, we first estimated the number of trips each vessel made in a month using the following formula:

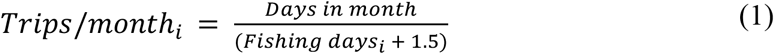

**Table 1.**
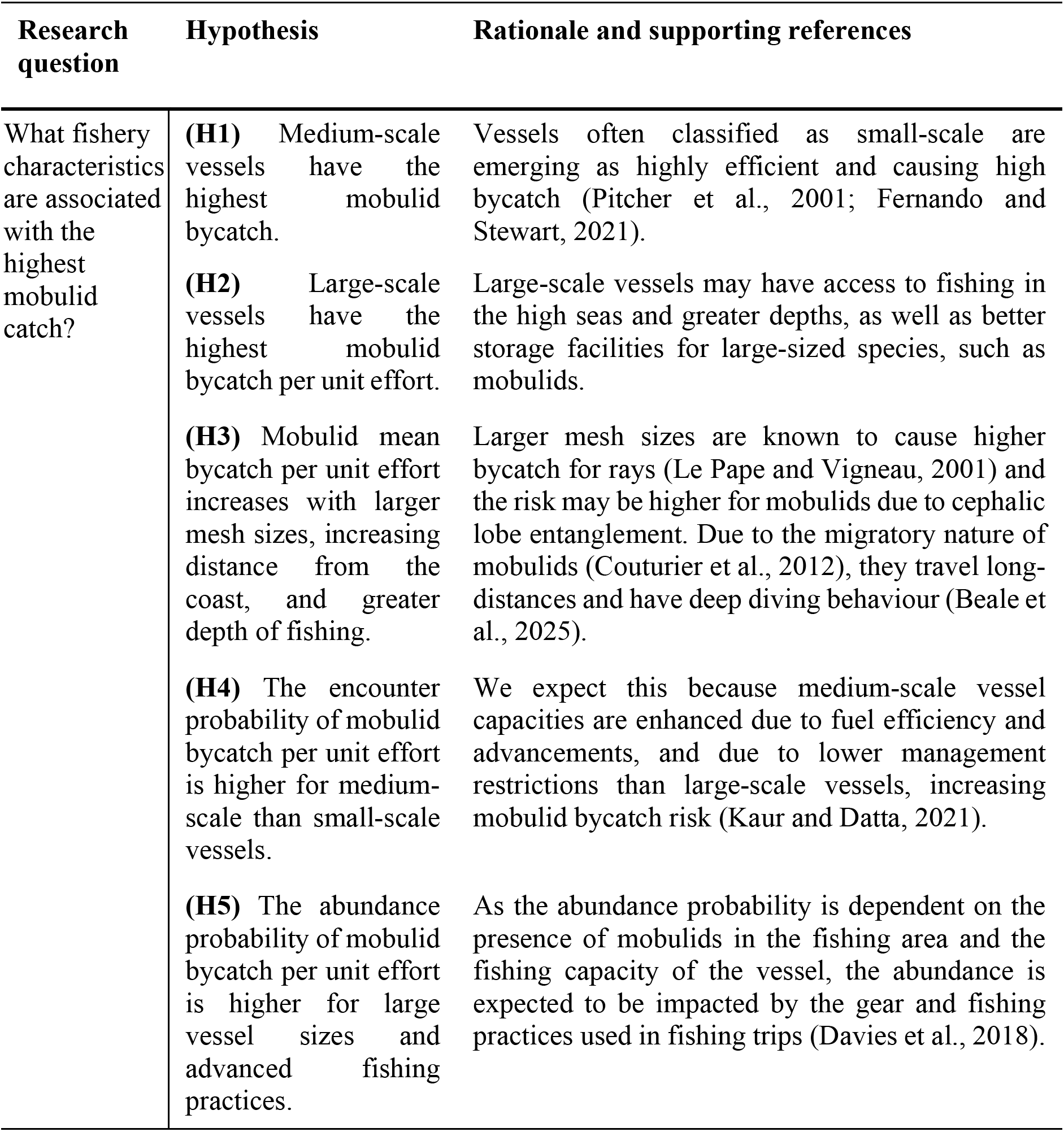
Hypotheses posed to identify fishery characteristics associated with mobulid bycatch landings in Thoothukudi District, Tamil Nadu, India.

| Research question | Hypothesis | Rationale and supporting references |
| --- | --- | --- |
| What fishery characteristics are associated with the highest mobulid catch? | (H1) Medium-scale vessels have the highest mobulid bycatch. | Vessels often classified as small-scale are emerging as highly efficient and causing high bycatch (Pitcher et al., 2001; Fernando and Stewart, 2021). |
|  | (H2) Large-scale vessels have the highest mobulid bycatch per unit effort. | Large-scale vessels may have access to fishing in the high seas and greater depths, as well as better storage facilities for large-sized species, such as mobulids. |
|  | (H3) Mobulid mean bycatch per unit effort increases with larger mesh sizes, increasing distance from the coast, and greater depth of fishing. | Larger mesh sizes are known to cause higher bycatch for rays (Le Pape and Vigneau, 2001) and the risk may be higher for mobulids due to cephalic lobe entanglement. Due to the migratory nature of mobulids (Couturier et al., 2012), they travel long-distances and have deep diving behaviour (Beale et al., 2025). |
|  | (H4) The encounter probability of mobulid bycatch per unit effort is higher for medium-scale than small-scale vessels. | We expect this because medium-scale vessel capacities are enhanced due to fuel efficiency and advancements, and due to lower management restrictions than large-scale vessels, increasing mobulid bycatch risk (Kaur and Datta, 2021). |
|  | (H5) The abundance probability of mobulid bycatch per unit effort is higher for large vessel sizes and advanced fishing practices. | As the abundance probability is dependent on the presence of mobulids in the fishing area and the fishing capacity of the vessel, the abundance is expected to be impacted by the gear and fishing practices used in fishing trips (Davies et al., 2018). |

Where, trips per month are estimated for an identified vessel *i*, which spends a number of fishing days to land the mobulids caught on the trip (Supplementary Information 1). A break period of 1.5 days was assumed between trips for restocking of food and storage ice, as per the personal communications with fisher community collaborators. The total trips per month were used to impute the zero mobulid bycatch trips for each vessel, and were scaled for the study period (April to October) (Supplementary Information 1). The positive count trips of mobulid catch per unit effort (CPUE) for each vessel were used to impute the zero mobulid bycatch trips for each vessel. The number of fishing days to calculate the total positive count trips was almost always taken as the median of the category range for data collected on ‘fishing days’ in the survey. The fishing day values assumed were: 1 day for category, ‘Single-day fishing’; 4 days for category ‘Multi-day: 2-6 days’; 9 days for category ‘Multi-day: 7-11 days’; 14 days for category ‘Multi-day: 12-16 days’; and 17 days for category ‘Multi-day: >16 days’. We recognise that a fixed number of rest days and fishing days is an assumed average, and there may variation in these values between trips. However, assuming an average is required to impute the number of trips for each vessel. We account for this potential variation by using a robust modelling approach and conducting sensitivity analysis. We only retained the boat size and mesh size information for the imputed trips as these were fixed assets, whereas other fishery characteristic variables (fishing depth, fishing distance, soak hours, fishing days) may vary between observed and unobserved trips. The unobserved and observed fishing trips of all identified vessels (with zero mobulid bycatch and positive mobulid CPUE) constituted the complete dataset.

#### 4.3.2 Multivariate analysis and Hurdle model

To assess multicollinearity among fishery characteristic variables, we constructed a Spearman correlation matrix suitable for ordered variables (Cotter, 2009) (Supplementary Information 2). The correlation value threshold of 0.6 was applied to reduce redundancy in parameter estimates (Artusi et al., 2002). As the site was a nominal variable, its correlation with ordered variables could not be meaningfully assessed in the correlation matrix; therefore, we examined site structure with covariates using a Principal Component Analysis (PCA) (Supplementary Information 2).

To model mobulid CPUE, we first assessed overdispersion and potential outlier contamination by estimating the dispersion ratio using the variance and mean of the mobulid CPUE data. To estimate the effect of fishery characteristics on mobulid CPUE, we selected a Hurdle model (Zeileis et al., 2008). Hurdle models are two component models with a truncated count component and a hurdle component that models zero versus positive counts. A hurdle model was the most suitable choice for our data, assuming there are two underlying processes determining presence or absence of mobulid catch and abundance of catch (Zeileis et al., 2008). We assume that two processes occur for each count value of CPUE: (1) encounter probability: We observe mobulid landings (num = 0/1) when they are absent or present in the fishing area (migrated or diving to non-fished depths) (2) abundance probability: The abundance of mobulids observed (num>0) depends on the presence or absence of mobulids (num = 0/1) and the efficiency of fishing practices.

Unlike traditional count models, such as the standard Poisson or Negative Binomial, which frequently fail to address zero inflation in large datasets, Hurdle models perform substantially better under conditions of zero inflation (Pitsha et al., 2025). To account for overdispersion, we specified a negative binomial distribution for the positive count component in the Hurdle model, given by (Pitsha et al. 2025):

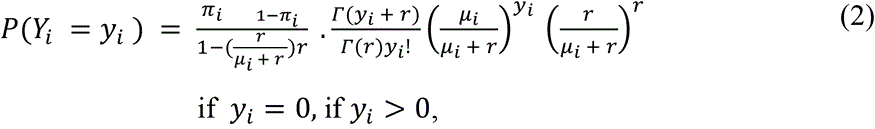

Where *Y_i_* is the count response for the i-th observation, and *i* = 1, …, *n*, and *n* is the total number of catch per unit effort observations. In (2), *r* is the dispersion parameter and *μ_i_* is the mean of the negative binomial distribution, *Γ* is the gamma function, and *π_i_* is the probability of observing a zero catch per unit effort. Fishery characteristic covariates were incorporated into both the zero and count components, as:

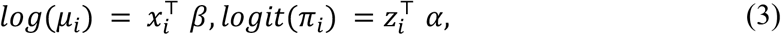

Where **α** and ***β*** are vectors of regression coefficients for the fishery characteristic covariates *x_i_* and *z_i_* respectively.

Although Hurdle negative binomial models effectively handle datasets with excess zeroes and low overdispersion, their performance becomes significantly poorer in the presence of zero inflation coupled with high overdispersion and outlier contamination (Pitsha et al., 2025). Marine megafauna bycatch data frequently contain extreme values (when an aggregation is caught) that can exert disproportionate influence on maximum likelihood estimates. To address overdispersion, zero inflation, and the sensitivity of the negative binomial Hurdle model to such influential outlier observations, we employed a Huber-robust extension to the hurdle model, with Huber’s ψ-function (Cantoni and Zedini, 2011). The Robust Hurdle Negative Binomial (RHNB) model iteratively reweights the maximum likelihood, downweighing only extreme residuals while maintaining the efficiency for the remainder of the data. The RHNB model is the most reliable option under substantial overdispersion and moderate outlier contamination, as it handles excess zeroes while retaining robustness against noise induced by extreme values (Pitsha et al., 2025). The RHNB model methodology implemented in this study follows Cantoni and Zedini (2011) and Pitsha et al. (2025), where a bounded influence is applied with cutoff points based on the mean and variance of catch per unit effort distribution. The RHNB model truncated negative binomial (NB) component can be expressed as:

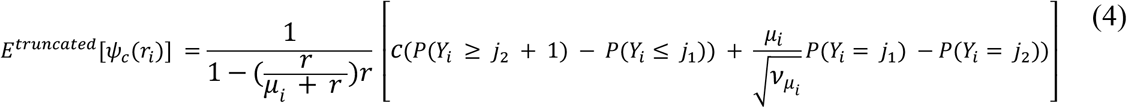

Where *Yi ∼ NB (μ_i_, r),* variance *v_μi_,* and cutoff points are, *j1* and *j2,*

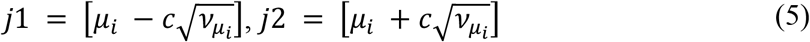

The constant *c* in the *ψ_c_* function tunes the balance between robustness and efficiency of the RHNB model. Usually, a standard *c* = 1.345 representing 95% efficiency is applied (Cantoni and Zedini, 2011). To confirm an appropriate Huber tuning constant suitable to our CPUE data in the RHNB model, we conducted a sensitivity analysis across alternative tuning constants to evaluate the stability of coefficient estimates (Supplementary Information 2). Once the tuning constant was selected, we ran the RHNB model using a maximum of 20 iterations until convergence, with a tolerance of 1e-6, assuming additional iterations would not meaningfully alter parameter estimates. Finally, we compared the standard Hurdle Negative Binomial model with the RHNB model using Akaike Information Criterion (AIC) and Bayesian Information Criterion (BIC) to evaluate model goodness of fit for our mobulid catch per unit effort data.

## 5. Results

### 5.1 Mobulid total catch and catch per unit effort

We report the mobulid total catch and mean catch per unit effort (per trip) for all fishery characteristic categories for the observed fishing trips (positive CPUE counts only) (Table 2; Figure 2). Over 99% of the mobulid landings were from gillnet fishing (as reported in Chopra et al., 2026a). The Tharuvaikulam Fishing Harbour fishing fleet had a higher total mobulid catch and CPUE than the Threspuram Fishing Village fleet despite a lower number of fishing trips. Seasonally, the month of August had the highest mobulid catch despite a similar number of fishing trips throughout June to September (Table 2).

**Figure 2.**
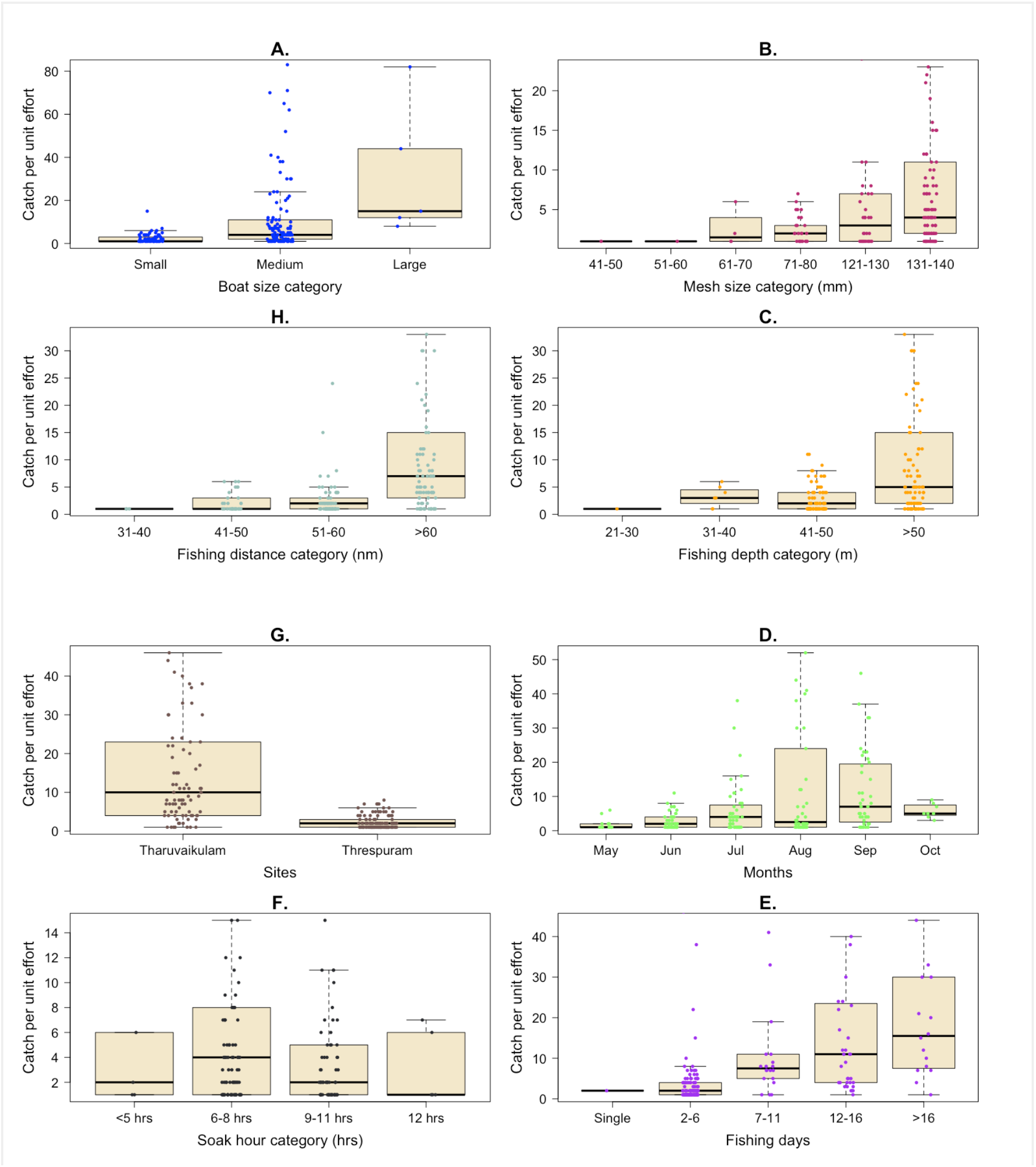
Catch per unit effort contributed from each fishery characteristic category in landings observed at study sites in Thoothukudi District, Tamil Nadu, India. The jittered points denote the catch per unit effort (per fishing trip). **A.** Boat sizes; **B.** Mesh sizes; **C.** Fishing depth; **D.** Months; **E.** Fishing days; **F.** Soak hours; **G.** Sites; **H.** Fishing distance.

**Table 2.** Fishery characteristic categories of the observed trips contributing to total mobulid catch at Threspuram Fishing Village (FV) and Tharuvaikulam Fishing Harbour (FH) in Thoothukudi District, Tamil Nadu, India.

| **Boat size (m)** |  |  |  |
| --- | --- | --- | --- |
| **Category** | **Mean CPUE**<br>**(individuals)** | **Total catch**<br>**(individuals)** | **Number of fishing**<br>**trips** |
| Small-scale (<15) | 2.29 | 156 | 68 |
| Medium-scale (15 – 21) | 10.90 | 1235 | 113 |
| Large-scale (>21) | 32.20 | 161 | 5 |
| Unidentified | - | - | 12 |
| **Mesh size (mm)** |  |  |  |
| **Category** | **Mean CPUE**<br>**(individuals)** | **Total catch**<br>**(individuals)** | **Number of fishing**<br>**trips** |
| 41-50 | 1 | 1 | 1 |
| 51-60 | 1 | 1 | 1 |
| 61-70 | 2.50 | 10 | 4 |
| 71-80 | 2.37 | 71 | 30 |
| 121-130 | 5.38 | 172 | 32 |
| 131-140 | 11.80 | 1238 | 105 |
| Unidentified | - | - | 25 |
| **Distance of fishing (nautical miles)** |  |  |  |
| **Category** | **Mean CPUE**<br>**(individuals)** | **Total catch**<br>**(individuals)** | **Number of fishing**<br>**trips** |
| 31-40 | 1 | 3 | 3 |
| 41-50 | 2.30 | 69 | 30 |
| 51-60 | 2.81 | 177 | 63 |
| > 61 | 14.30 | 1285 | 90 |
| Unidentified | - | - | 12 |
| **Depth of fishing (m)** |  |  |  |
| **Category** | **Mean CPUE**<br>**(individuals)** | **Total catch**<br>**(individuals)** | **Number of fishing**<br>**trips** |
| 21-30 | 1 | 1 | 1 |
| 31-40 | 3.29 | 23 | 7 |
| 41-50 | 2.82 | 220 | 78 |
| >50 | 12.80 | 1269 | 99 |
| Unidentified | - | - | 13 |
| **Site** |  |  |  |
| **Category** | **Mean CPUE**<br>**(individuals)** | **Total catch**<br>**(individuals)** | **Number of fishing**<br>**trips** |
| Threspuram FV | 2.29 | 259 | 113 |
| Tharuvaikulam FH | 17.80 | 1509 | 85 |
| **Month** |  |  |  |
| **Category** | **Mean CPUE**<br>**(individuals)** | **Total catch**<br>**(individuals)** | **Number of fishing**<br>**trips** |
| May | 2.09 | 23 | 11 |
| June | 2.88 | 121 | 42 |
| July | 8.45 | 431 | 51 |
| August | 15.0 | 692 | 46 |
| September | 11.40 | 455 | 40 |
| October | 5.75 | 46 | 8 |

**Soak hours**
| **Category** | **Mean CPUE**<br>**(individuals)** | **Total catch**<br>**(individuals)** | **Number of fishing**<br>**trips** |
| --- | --- | --- | --- |
| = <5 | 18.40 | 92 | 5 |
| 6-8 | 8.82 | 811 | 92 |
| 9-11 | 4.57 | 265 | 58 |
| 12-14 | 3.20 | 16 | 5 |

**Days spent fishing**
| **Category** | **Mean CPUE**<br>**(individuals)** | **Total catch**<br>**(individuals)** | **Number of fishing**<br>**trips** |
| --- | --- | --- | --- |
| Single day | 2 | 2 | 1 |
| 2-6 days | 3.99 | 479 | 120 |
| 7-11 days | 12.20 | 245 | 20 |
| 12-16 days | 18.20 | 638 | 35 |
| >16 days | 20.50 | 328 | 16 |
| Unidentified | - | - | 6 |

Our results support H1–H2 (Table 1), indicating that medium-scale vessels have the highest total mobulid bycatch (80% of mobulid bycatch) and large-scale vessels had the highest bycatch per unit effort of 32.20 individuals, although this was only contributed by five fishing vessels (Table 2; Figure 3). We recorded 63 small-scale (37%), 101 medium-scale (60%), five large-scale vessels (3%), and 12 unidentified size-class vessels (7%), based on our vessel size classification definitions (Table 2). Our results support H3, which states that mobulid mean bycatch per unit effort increases with larger mesh sizes, increasing distance from coast, and greater depth of fishing (Table 2; Figure 3). The mobulid catch increased monotonically with ordered categories of mesh size, fishing distance, and fishing depth. The largest recorded mesh size of 131-140 mm contributed 83% of the mobulid catch; vessels fishing >60 nautical miles away resulted in 84% of the mobulid catch; and fishing at depth over 50 m contributed to 84% of the mobulid catch (Figure 3).

**Figure 3.**
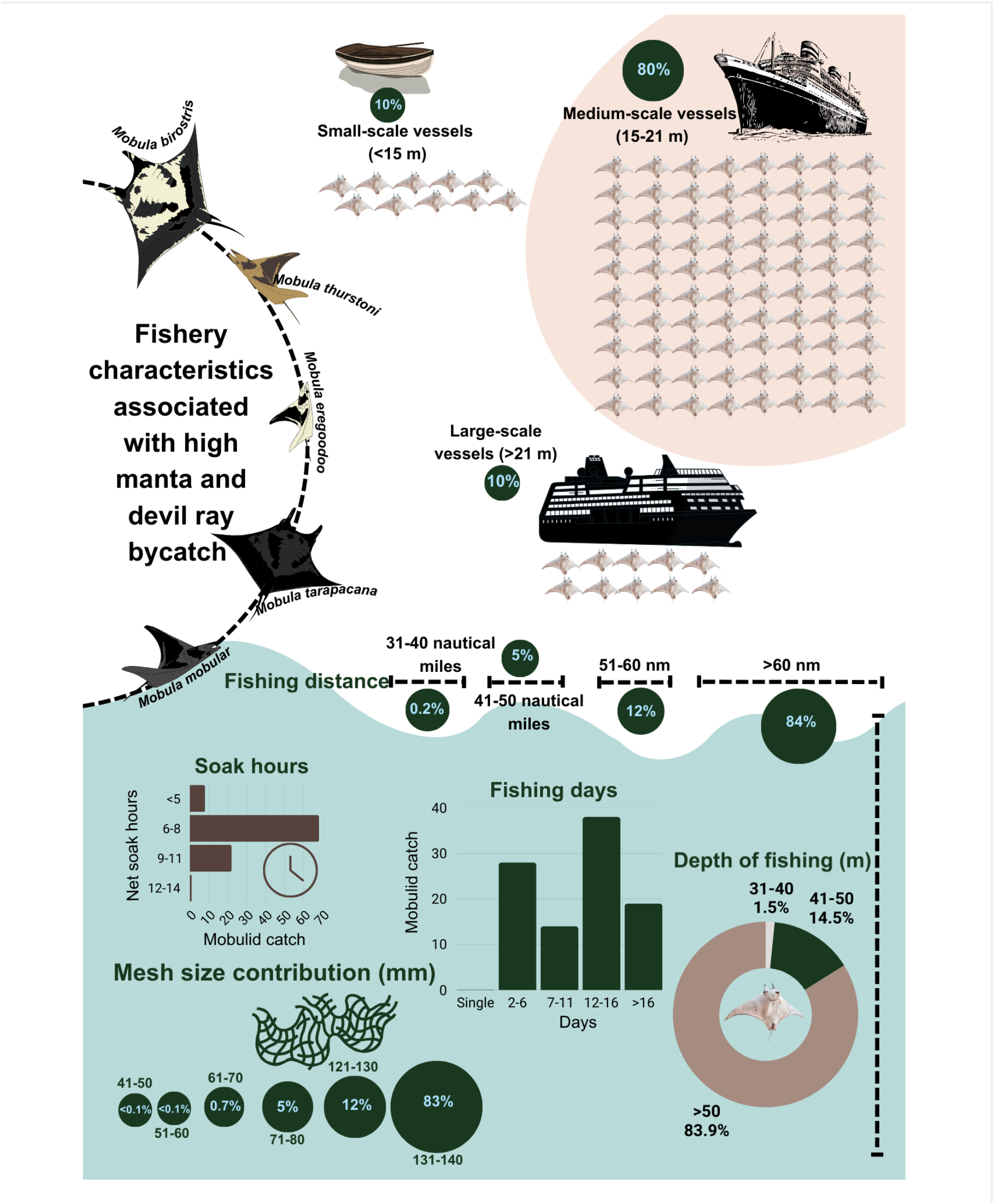
Fishery characteristics associated with observed mobulid landings at study sites in Thoothukudi District, Tamil Nadu, India. Study sites include Threspuram Fishing Village and Tharuvaikulam Fishing Harbour, Tamil Nadu, India for surveys conducted between April to October 2023. The percent values represent the associated catch contribution of the fishery characteristic variable to total mobulid catch.

### 5.2 Multivariate analysis and Hurdle model

The Spearman correlation matrix between fishery characteristic variables showed low to moderate correlation coefficients below the threshold of 0.6 (Supplementary Information 2). Therefore, we retained all fishery characteristic variables for the RHNB model, including boat size, mesh size, fishing distance, fishing depth, soak hours, and fishing days. Principal component analysis (PCA) showed that major differences in the created orthogonal principal components were correlated with site differences (Supplementary Table 2.2). The PCA was useful in deciphering the nature of collinearity among fishery characteristics and confirmed that a major factor for correlations was due to Threspuram Fishing Village and Tharuvaikulam Fishing Harbour fleets exhibiting different fishing practices. To avoid masking the impact of fishery characteristics on mobulid catch per unit effort due to the site variable, we did not include site in the RHNB model.

Our zero inflated data had high overdispersion and moderate outlier contamination. The variance of our distribution was significantly higher than the mean, and the dispersion ratio was 33.46. The distribution of positive count data showed upper-tail outliers, as extreme CPUE values were likely being observed when large mobulid aggregations were caught (Supplementary Information 2). Over 95% of the positive CPUE observations were under 40.15, and 99% were under 71.33, and the influential upper-tail outliers occurred in less than 5% of the CPUE observations. Thus, high overdispersion and moderate outlier contamination justified our choice of using the Huber-robust extension to a Hurdle negative binomial model. Sensitivity checks using alternative Huber tuning constant (*c*) produced stable coefficient estimates for values between 1.1–1.5 (Supplementary Information 2), supporting our choice of applying the standard *c* = 1.345, representing 95% data usage efficiency. The RHNB model (AIC: 850, BIC: 934) was a significantly better fit for our mobulid CPUE data, than the standard Hurdle negative binomial model (AIC: 2099, BIC: 2184).

Our results support H4, stating that the encounter probability of mobulid catch per unit effort is higher for medium-scale than small-scale vessels (Table 3). Medium-scale vessels are nearly twice as likely to catch a mobulid (non-zero CPUE probability) than small-scale vessels, while large-scale vessels are about six times more likely to record a non-zero mobulid CPUE (Table 3). Additionally, a larger mesh size also significantly increases the chances of a non-zero mobulid CPUE (Table 3). Our results support H5, which states that the abundance probability of mobulid catch per unit effort is higher for intensive fishing practices (Table 3). A higher number of days spent fishing resulted in a significantly higher abundance of catch per unit effort, representing fishing intensity. Every extra five days spent fishing increases the expected mobulid CPUE abundance approximately by two-fold (Table 3). However, other fishery characteristics, such as boat size, mesh size, fishing depth, fishing distance, and soak hours did not show a significant impact on the abundance probability of mobulid catch per unit effort. Notably, in the standard hurdle negative binomial model (Supplementary Information 2), medium and large-scale vessels exhibit a weak positive association with CPUE abundance (p-value <0.1), and fishing distance shows a positive association with CPUE abundance (p-value <0.05). However, this effect is not observed in the RHNB model, as expected, because outliers (extreme CPUE counts) are down-weighted in maximum likelihood estimation.

**Table 3.** Model estimates of Huber-robust Hurdle Negative binomial model for catch per unit effort per fishing trip of mobulid bycatch in Thoothukudi District, Tamil Nadu, India. For the count part, the positive catch per unit effort is the response variable and fishery characteristics (boat size, mesh size, fishing depth, fishing distance, and fishing days) as independent variables. For the zero part, boat size and mesh size are independent variables. Note: Significance indicated by asterisks p < 0.05 (*), p < 0.01 (**), p < 0.001 (***).

| **Count model coefficients (truncated negative binomial with log link)** |  |  |  |  |  |
| --- | --- | --- | --- | --- | --- |
|  | **Estimate** | **Odds ratio vs reference** | **Std. error** | **z-value** | **p-value** |
| Intercept | -7.496 | - | 3.332 | -2.250 | < 0.05 * |
| Boat size (Medium-scale) | 0.599 | 1.82x higher odds | 0.594 | 1.010 | 0.312 |
| Boat size (Large-scale) | 1.643 | 5.17x higher odds | 1.048 | 1.568 | 0.116 |
| Mesh size | 0.076 | 1.08x higher odds | 0.176 | 0.435 | 0.663 |
| Fishing depth | -0.043 | 0.96x lower odds/unit increase | 0.482 | -0.090 | 0.927 |
| Fishing distance | 0.650 | 1.92x high odds/unit increase | 0.411 | 1.581 | 0.113 |
| Soak hours | 0.410 | 1.51x high odds/unit increase | 0.458 | 0.896 | 0.370 |
| Fishing days | 0.698 | 2.01x high odds/unit increase | 0.257 | 2.708 | < 0.01 ** |
| Log (theta) | -0.532 | - | 0.635 | -0.838 | 0.402 |
| **Zero hurdle model coefficients (binomial with logit link)** |  |  |  |  |  |
|  | **Estimate** | **Odds ratio vs reference** | **Std. error** | **z value** | **p-value** |
| Intercept | -6.011 | - | 0.576 | -10.422 | < 0.001 *** |
| Boat size (Medium-scale) | 0.671 | 1.96x higher odds | 0.300 | 2.238 | < 0.05 * |
| Boat size (Large-scale) | 1.787 | 5.97x higher odds | 0.626 | 2.855 | < 0.01 ** |
| Mesh size | 0.133 | 1.14x higher odds/unit increase | 0.061 | 2.157 | < 0.05 * |

| **Pearson residuals** |  |  |  |  |
| --- | --- | --- | --- | --- |
| **Minimum** | **First Quantile** | **Medium** | **Third Quantile** | **Maximum** |
| -0.234 | -0.134 | -0.096 | -0.069 | 17.180 |

## 6. Discussion

Using mobulid landings and fishery data, we inform fishery management prioritisation for manta and devil ray conservation. Our results provide evidence that fishing practices significantly impact the presence and abundance of mobulids in landed catch, supporting H1– H5. We find that medium-scale vessels are the largest contributors to manta and devil ray landings in our case study (H1), while large-scale vessels show the highest mobulid catch per unit effort (H2). Mobulid mean catch per unit effort increases with larger mesh sizes, increasing distance from the coast, and greater depth of fishing (H3). Using the Huber-robust extension to the Hurdle negative binomial model, we find that medium-scale vessels are twice as likely to land a mobulid, in support of H4, meaning the mobulid encounter probability of medium-scale vessels is significantly higher than that of small-sized vessels. Finally, we find that higher fishing intensity (denoted by fishing days), significantly increases the probability of catching a higher number of manta and devil rays per trip by two-fold. We propose a more nuanced fisheries management approach, departing from the binary definitions of small-scale and large-scale fisheries, which is especially essential for applying appropriate regulatory frameworks and support for fishing communities and for sustainable fisheries development (Perez et al., 2012; Tseng and Kao, 2022). With a focus on marine megafauna conservation, we show through our case study that the probability of medium-scale vessels catching a mobulid is twice that of small-scale vessels, and that the abundance of mobulid catch increases with intensified fishing duration. The ability to engage in intensive fishing practices, resulting in a significantly higher encounter and abundance bycatch probability, indicate capacity differences, which have been observed as an emerging threat to vulnerable megafauna and individual fisheries sustainability globally (Alfaro-Shigueto et al., 2010; da Silva et al., 2023; Saüt et al., 2024).

We recognise that the classification of fisheries capacities is complex and includes multiple fisheries characteristics in addition to vessel length or size, as proposed through a vast body of historical literature examining fisheries classifications (Smith and Basurto, 2019). However, about 51–65% of the scientific literature accounts for boat size, length, or capacity as a starting point to examine fishery classifications (Chuenpagdee et al., 2006; Smith and Basurto, 2019). Beyond vessel length, fisheries value chain dynamics (pre-harvest, harvest, and post-harvest) have also been proposed as valuable metrics for SSF classification (Smith and Basurto, 2019). Here we emphasise the contribution of medium-scale vessels. Our aim is not solely to re-classify fisheries based on vessel size, but rather twofold: first, to clarify the definition of SSF in India to enable effective SSF management and governance; and second, to make a capacity distinction within the globally classified SSFs. This distinction includes a low-capacity subsistence class and an emerging medium-capacity class of fisheries that requires a higher degree of restrictive management measures for conservation (as evidenced by our results). We want to emphasise that the recommendation to depart from binary classification is not divergent from, but rather a complementary measure to, FAO’s “SSF characterisation matrix” (Funge-Smith, 2018). Briefly, the SSF characterisation tool scores fisheries (on a matrix) based on a range of qualitative indicators that can be used to assess the scale of SSF nationally when compared globally (Funge-Smith, 2018). The cutoff point between small- and large-scale is then context-dependent within the country. We believe that the characterisation matrix is a positive development toward better-managed fisheries; however, in practice, if vessels are reclassified into binary categories of small and large after scoring, the national fisheries management regulations are applied in two tiers. We therefore argue that, in practice, at this cutoff stage, there should be a three-tier classification of small, medium, and large, which is long overdue and especially relevant for rapidly expanding blue-economy nations such as India, because the regulation requirements between the three tiers are too vast to be merged into two classifications. We also note that, we do not solely fixate on technological differences between fisheries; there are likely socio-economic and ownership dynamics that differ between the aforementioned small- and medium-scale vessels (Jadhav, 2018); however, exploring these distinctions falls outside the scope of this study.

Poor management measures enable overfishing, leading to dwindling stocks of target and bycatch species and global fisheries unsustainability (Sharma et al., 2025). Vessel size classifications and fisheries policies are often incongruent in national laws; for example, the Indian Marine Policy (India G.O., 2004) defines marine fish resources classified into subsistence fishing (<12 m), small-scale fishing (12-20 m) and large-scale fishing (>20 m); whereas the Indian Marine Fisheries Act (India G.O., 2021) applies regulations to vessels as <15 m or >15 m overall length for penalties (applied to both mechanised and motorised vessels). Additionally, there are varying measures that aid in state fisheries management through state-specific Marine Fisheries Regulation Acts (Sahu et al., 2022). While there are multiple policy frameworks and general recognition of coastal overcapacity in Indian fisheries (Gangal et al., 2023), there remains: (1) a lack of biologically relevant fisheries management (Gangal et al., 2023), (2) a lack of size limit bans for elasmobranchs, despite evidence of age-class truncation in literature (Kopf et al., 2025; Chopra et al., 2026a), and (3) current mesh and minimal landing size for species regulations not tied to vessel capacities across states (Gangal et al., 2023). Such inconsistencies in vessel size regulations, and subsequent mesh-size and minimal landing size fisheries regulations associated with those classifications, endanger species viability of elasmobranchs in both SSF and LSF. In this study, we classify small-scale vessels as <15 m in length and find differences in mobulid total catch and CPUE contributions among vessel size categories. Consequently, we recommend that fisheries management measures distinguish vessel capacities also in terms of elasmobranch bycatch risk to apply associated biologically-relevant regulations. We argue that failure in distinguishing vessel capacities poses risks in the regulatory and welfare contexts. In the regulatory context, there is a risk of applying stringent regulations to subsistence fisheries, or inversely applying diluted measures to advanced medium-capacity fisheries (where stricter regulations are warranted). In the welfare context, it risks the division of subsidiary resources which are intended for supporting subsistence class fishers (Harper and Sumaila, 2019).

Prioritising fisheries management based on fleet characteristics streamlines policy implementation, thereby increasing the efficiency of conservation measures. For example, in our case study, we emphasise the importance of high capacity medium-scale vessels as a priority sector for conservation measures for manta and devil rays. We also recommend prioritisation of fisheries that use larger mesh sizes for manta and devil ray conservation, supported by a significantly higher contribution of large mesh sizes in our results. Mesh size specificity in bycatch has been identified as a potential issue, particularly for elasmobranchs with protruding head shapes (cephalic lobes for manta and devil rays) causing a higher degree of net entanglement. This is evident in, for example, hammerhead sharks (Thorpe and Frierson, 2009) and sawfish (Dulvy et al., 2016). The entanglement effect of the net, and thereby bycatch, also increases by reducing the hang due to minimised buoyancy (Northridge et al., 2017), which can likely occur when non-standard buoyancy devices are used. Indirect incentive-based policy (For example, net repair subsidies) instruments (Booth et al., 2025) could support release policies for large mesh sizes (>100 mm), which are predominantly used in tuna-targeted fisheries (Squires and Garcia, 2018) and account for the mobulid bycatch observed in this study. However, incentive-based approaches may have variable effects on both bycatch and target species, making research trials and attention to design complexity key in avoiding unintended consequences and counter-productive fisher behaviour (Finkelstein et al., 2008; Booth et al., 2025).

In conclusion, we propose that within the globally classified small-scale fisheries, medium-scale vessels with advanced capacities and efficiency are an emerging problem for fisheries management. Our results support a comprehensive re-conceptualisation of small-scale fisheries complementary to FAO’s SSF characterisation matrix (Fungee-Smith, 2018), such that advanced medium-capacity vessels are subject to fisheries management measures commensurate with their bycatch impact on elasmobranchs. The re-conceptualisation could help address challenges in reducing their impact, support bycatch mitigation of vulnerable marine megafauna, promote sustainable fisheries development in emerging blue economy nations, and helping prevent local extinctions of endangered fauna such as manta and devil rays (Pardo et al., 2016). Therefore, we recommend aligning elasmobranch demographic considerations with management measures by applying biologically relevant regulations to vessel capacities that pose the highest risk of elasmobranch bycatch.

## 7. Acknowledgements

We thank our field assistants A. Nirojan, J. Dyson, and Ramesh in Tamil Nadu for their invaluable help with data collection. We are especially grateful to the fishers at the landing centres of Thoothukudi, who cooperated when the mobulid landing data was being collected. We would also like to thank the useful discussions with Sarah Bull which helped develop the ideas in this study. M.C. received funding through the Swami Vivekananda Scholarship for Academic Excellence (formerly Rajiv Gandhi Scholarship), granted by the State Government of Rajasthan, Government of India. Fieldwork and data collection were funded by a 2023 Graduate Student Research Award from the Society for Conservation Biology and the Emergency Grant by the Manta Trust.

