## Supplementary Information for "The emerging medium-scale: Prioritising fisheries management for elasmobranch conservation"

### 1. Data cleaning and imputation

As the vessel size data was collected using fisher responses, we found respondent variation on reported vessel size in approximately 5.9% of the observed vessels (n = 10 vessels and 27 trips) (Supplementary Table 1.1). This recorded variation likely occurred in cases where vessel size falls at the borderline between defined vessel size categories (small, medium, large). To ensure consistent handling of borderline cases with respondent variation, we implemented the following decision rules consistently across all observations:

- **Case I: Borderline between small-scale and medium-scale vessels**

In borderline cases between small-scale and medium-scale vessels (n = 6 vessels and 15 trips), we classified categories based on their site. As all the borderline cases between small-scale and medium-scale belonged to Threspuram Fishing harbour, we assigned these six vessels the size category dominant in Threspuram. This decision was informed by the overall distribution observed in the dataset (Supplementary Table 1.2), wherein 61% of vessels in Threspuram were small-scale and hence these 6 vessels had a higher probability of being small-scale.

- **Case II: Borderline between medium-scale and large-scale vessels**

One borderline case (n = 1), comprising two trips (Supplementary Table 1.1), occurred between the medium-scale and large-scale categories. As there was an equal likelihood that the vessel could be classified as either medium-scale or large-scale, we retained the first recorded value (Trip ID 103), which classified the vessel as medium-scale. This decision was also informed by the broader structure of our dataset, with very few large-scale vessels (n = 5) and a dominant proportion of medium-scale vessels (n = 101), therefore there's a higher probability that this vessel was a medium-scale vessel.

- **Case III: Borderline between small-scale and medium-scale vessels overlapping across sites**

Three borderline cases (n = 3) involved vessels operating across both Threspuram Fishing Village and Tharuvaikulam Fishing Harbour, with respondent variation in reported vessel size. Given that these vessels were at the borderline between size categories, and all other factors being equal, differences in fishing grounds between the two sites could reasonably influence mesh size or other fishery characteristics. For these cases, we utilised other vessel characteristics to determine appropriate size classification. For example, vessels described with smaller mesh sizes were retained as small-scale, while those with larger mesh sizes were retained as medium-scale to maintain internal consistency between fishery characteristics.

**Supplementary Table 1.1:** Vessels showing respondent variation in reported vessel size (n = 10) for mobulid landings in Thoothukudi District landing centres in southeast India in 2023 (Threspuram Fishing Village [TP] and Tharuvaikulam Fishing Harbour [TVK]). Vessel ID denotes the unique identification number assigned to each vessel in the dataset. Trip ID refers to the unique identifier assigned to each fishing trip undertaken by a vessel. The 'Assigned' column indicates the final vessel size category allocated to each Vessel ID, based on the consistent decision rules applied across all cases of respondent variation in reported vessel size.

| Vessel ID and cases | Trip ID and total mobulid catch (individuals) in each trip | Reported vessel sizes | Assigned |
|---|---|---|---|
| 5<br>(Case I) | 3 fishing trips:<br>Trip ID 5: 6<br>Trip ID 7: 5<br>Trip ID 36: 1 | Trip ID 5: Small-scale<br>Trip ID 7: Small-scale<br>Trip ID 36: Medium-scale | Small-scale |
| 20<br>(Case I) | 2 fishing trips:<br>Trip ID 22: 3<br>Trip ID 144: 1 | Trip ID 22: Small-scale<br>Trip ID 144: Medium-scale | Small-scale |
| 24<br>(Case I) | 2 fishing trips:<br>Trip ID 26: 1<br>Trip ID 129: 1 | Trip ID 26: Small-scale<br>Trip ID 129: Medium-scale | Small-scale |

| | | | |
|---|---|---|---|
| 37<br>(Case I) | 3 fishing trips:<br>Trip ID 41: 1<br>Trip ID 51: 1<br>Trip ID 154: 2 | Trip ID 41: Small-scale<br>Trip ID 51: Medium-scale<br>Trip ID 154: Medium-scale | Small-scale |
| 64<br>(Case I) | 2 fishing trips:<br>Trip ID 70: 2<br>Trip ID 192: 5 | Trip ID 70: Small-scale<br>Trip ID 192: Medium-scale | Small-scale |
| 98<br>(Case I) | 3 fishing trips:<br>Trip ID 106: 2<br>Trip ID 123: 1<br>Trip ID 138: 2 | Trip ID 106: Small-scale<br>Trip ID 123: Medium-scale<br>Trip ID 138: Medium-scale | Small-scale |
| 95<br>(Case II) | 2 fishing trips:<br>Trip ID 103: 8<br>Trip ID 119: 15 | Trip ID 103: Medium-scale<br>Trip ID 119: Large-scale | Medium-scale |
| 1<br>(Case III) | 3 fishing trips:<br>Trip ID 1: 1<br>Trip ID 110: 1<br>Trip ID 190: 11 | Trip ID 1: Small-scale<br>Trip ID 110: Small-scale<br>Trip ID 190: Medium-scale | TVK: Medium-scale<br>TP: Small-scale |
| 7<br>(Case III) | 5 fishing trips:<br>Trip ID 8: 2<br>Trip ID 29: 1<br>Trip ID 131: 2<br>Trip ID 170: 23<br>Trip ID 185: 22 | Trip ID 8: Small-scale<br>Trip ID 29: Medium-scale<br>Trip ID 131: Medium-scale<br>Trip ID 170: Medium-scale<br>Trip ID 185: NA | TVK: Medium-scale<br>TP: Small-scale |
| 18<br>(Case III) | 2 fishing trips:<br>Trip ID 20: 5<br>Trip ID 109: 12 | Trip ID 20: Small-scale<br>Trip ID 109: Medium-scale | TVK: Medium-scale<br>TP: Small-scale |

**Supplementary Table 1.2:** Overall distribution of vessel size categories in the fisheries dataset for mobulid landings in Thoothukudi District landing centres in southeast India in 2023. The table represents the vessel sizes dominant in each site that were used to allocate vessel size categories in cases of respondent variation (n = 6 vessels).

| **Site** | **Vessel size category** | **Number of vessels** | **Percent within each site (%)** |
|---|---|---|---|
| Tharuvaikulam | Small-scale | 3 | 3.57 |
| Tharuvaikulam | Medium-scale | 66 | 78.6 |
| Tharuvaikulam | Large-scale | 5 | 5.95 |
| Tharuvaikulam | Unidentified | 10 | 11.9 |
| Threspuram | Small-scale | 61 | 61 |
| Threspuram | Medium-scale | 37 | 37 |
| Threspuram | Unidentified | 2 | 2 |

**Supplementary Table 1.3** Positive count trips recorded for mobulid landings in Thoothukudi District landing centres in southeast India in 2023 (Threspuram Fishing Village [FV] and Tharuvaikulam Fishing Harbour [FH]).

| Site | Month | Trips recorded |
|---|---|---|
| Threspuram FV | April | 0 |
| | May | 11 |
| | June | 35 |
| | July | 30 |
| | August | 22 |
| | Sept | 9 |
| | Oct | 6 |
| Tharuvaikulam FH | April | 0 |
| | May | 0 |
| | June | 7 |
| | July | 21 |
| | August | 24 |
| | Sept | 31 |
| | Oct | 2 |

### 2. Multivariate analysis and Hurdle model supplementary

The Spearman correlation matrix showed low to moderate multicollinearity between the ordered fishery characteristic variables of boat size, mesh size, fishing distance, fishing depth, net soak time, and fishing days (Supplementary Figure 2.1).

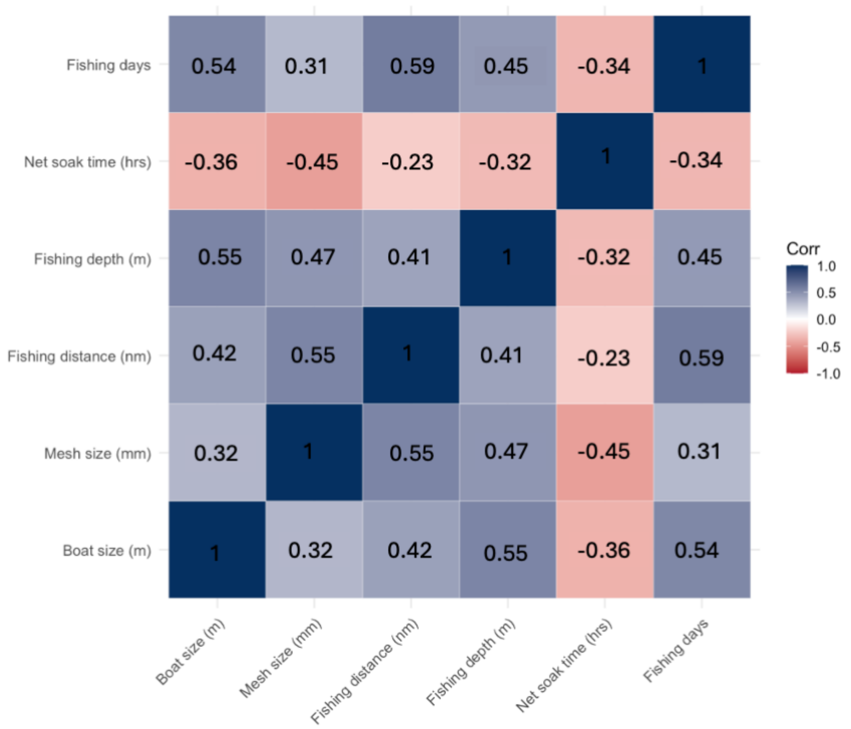


**Supplementary Figure 2.1** Spearman correlation matrix between fishery characteristic variables, boat size, mesh size, distance of fishing from coast, soak hours and, days spent fishing for mobulid bycatch monitored in 2023 at Threspuram Fishing Village and Tharuvaikulam Fishing Harbour in Tamil Nadu, India.

To understand the multicollinearity between variables with respect to site differences, we conducted a Principal Component Analysis (PCA). Principal component analysis is used when the correlation between components is moderate to high, with an aim to explain variance by creating uncorrelated orthogonal variables called principal components, which reduce dimensionality and mitigate information loss (Abdi and Williams, 2010; Jolliffe and Cadima, 2016). Additionally, PCA creates new variables that are linear combinations of fishery characteristic variables (Abdi and Williams, 2010), which is useful in our study for identifying the occurrence of site clustering due to different fishing practices in each site. As our fishery characteristic categories were coded numeric, ordered with multiple level categories (similar to continuous variables), the distance between categories was one point and therefore a numeric PCA was applied (Supplementary Table 2.1; Supplementary Figure 2.2). The fishery principal components PC1–PC3 explain the cumulative proportion of over 75% of variation in catch per unit fishing trip (Supplementary Table 2.1; Supplementary Figure 2.2). We show fishery characteristic combinations in Threspuram Fishing Village and Tharuvaikulam Fishing Harbour for PC 2 and PC 3 with PC1 (Supplementary Figure 2.3).

**Supplementary Table 2.1** Cumulative proportion of variance in mobulid catch per unit effort, explained by principal components of fishery characteristic variables for mobulid bycatch monitored in 2023 at Threspuram Fishing Village and Tharuvaikulam Fishing Harbour in Tamil Nadu, India.

| | **PC1** | **PC2** | **PC3** | **PC4** | **PC5** | **PC6** |
|---|---|---|---|---|---|---|
| **Standard deviation** | 1.764 | 0.958 | 0.846 | 0.730 | 0.668 | 0.520 |
| **Proportion of Variance** | 0.519 | 0.153 | 0.119 | 0.088 | 0.074 | 0.045 |
| **Cumulative proportion** | 0.519 | 0.672 | 0.791 | 0.880 | 0.954 | 1.000 |

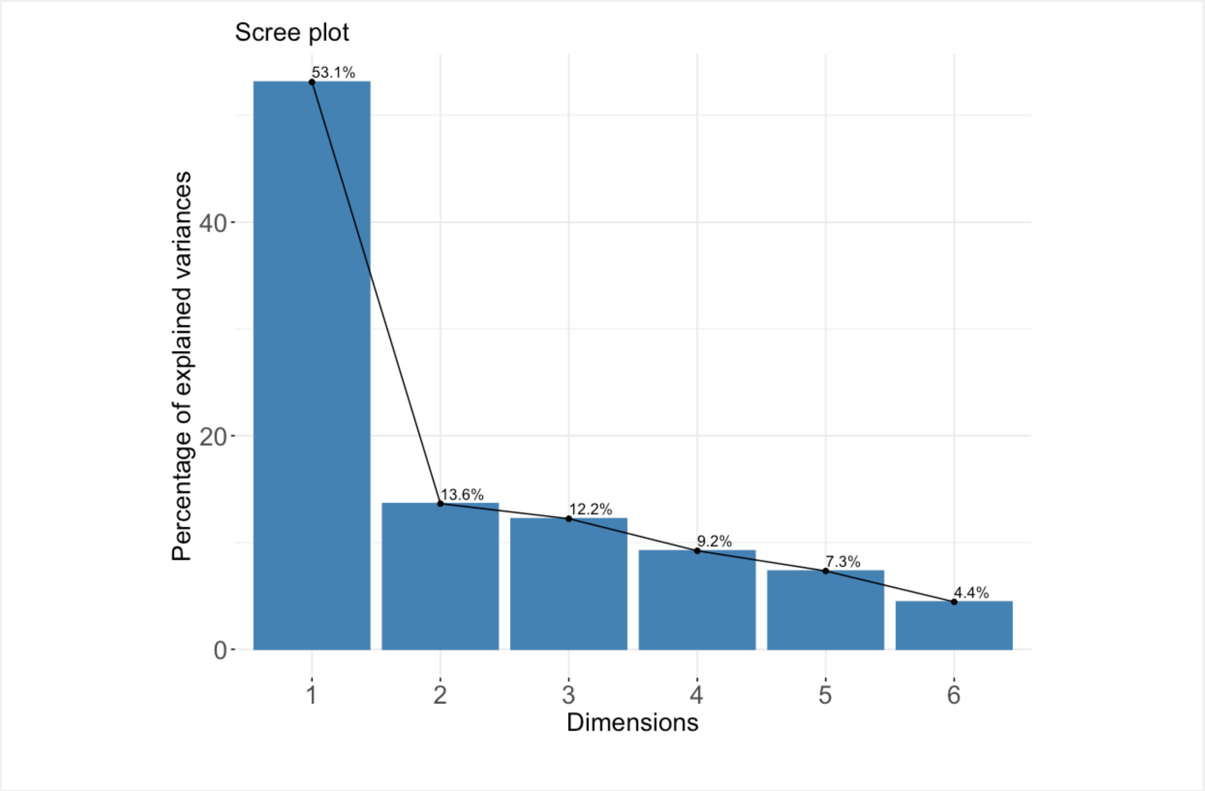

**Supplementary Figure 2.2** Proportion of variance in mobulid catch per unit effort explained by principal component dimensions of variables boat size, mesh size, distance of fishing from coast, soak hours and, days spent fishing for mobulid bycatch monitored in 2023 at Threspuram Fishing Village and Tharuvaikulam Fishing Harbour in Tamil Nadu, India.

79

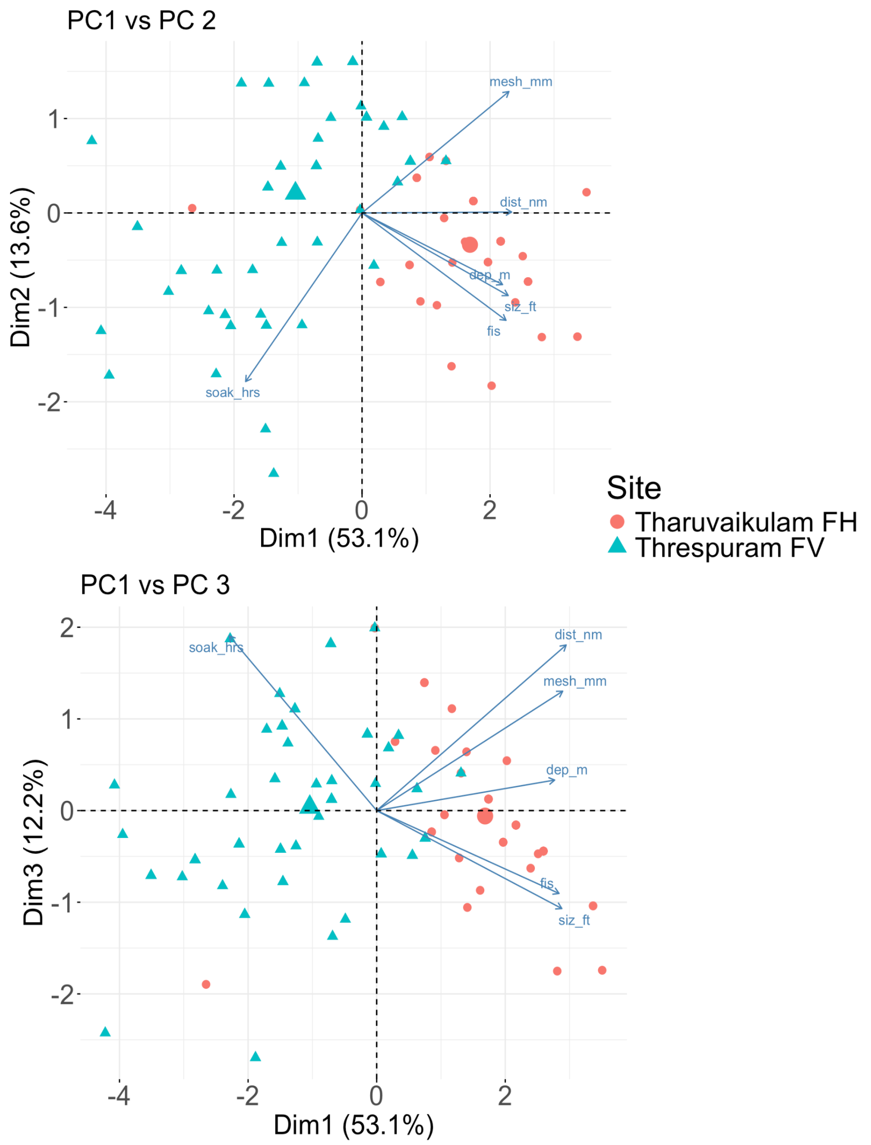


**Supplementary Figure 2.3** Site structure and clustering in fishery characteristic PCA dimensions (PC1–PC3) showing differences in fishery practices between sites Threspuram Fishing Village (FV) and Tharuvaikulam Fishing Harbour (FH). The principal component dimensions constitute fishery characteristic variables, boat size, mesh size, distance of fishing from coast, depth of fishing, soak hours, and days spent fishing for mobulid bycatch observed in 2023 at Threspuram Fishing Village and Tharuvaikulam Fishing Harbour in Tamil Nadu, India.

80

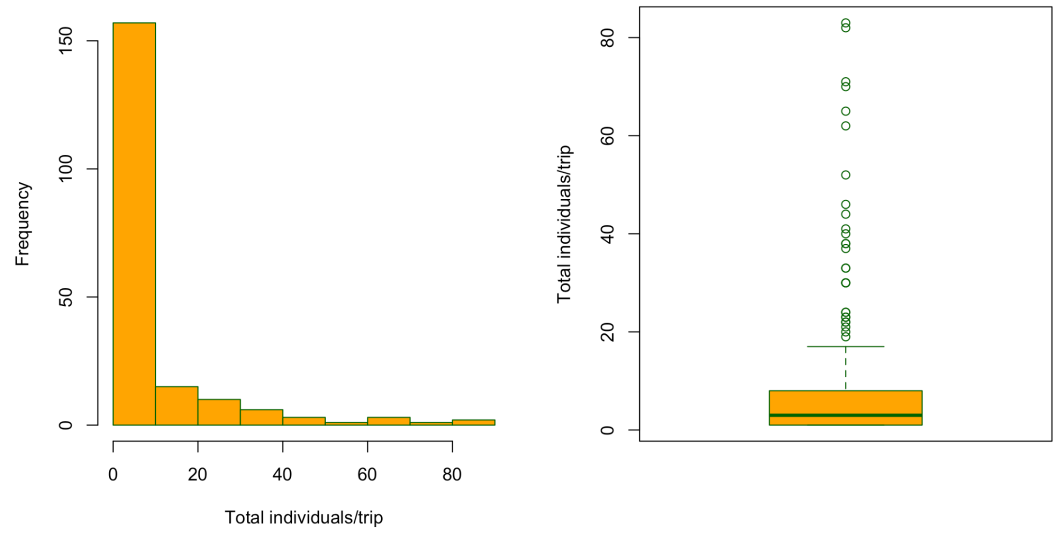


**Supplementary Figure 2.4** Outlier contamination and overdispersion in mobulid catch per unit effort (per fishing trip) in observed landings in 2023 in Tamil Nadu, India.

81

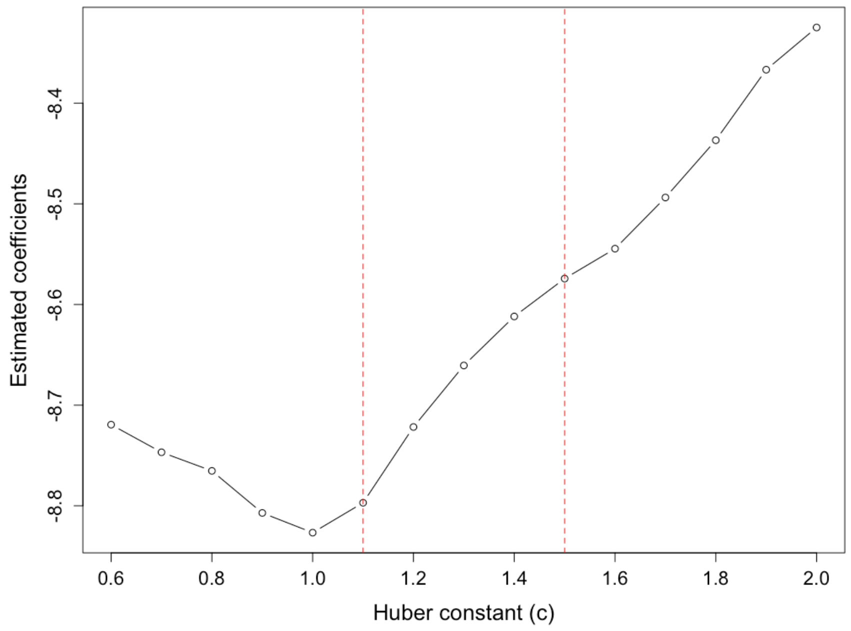


**Supplementary Figure 2.5** Plot showing the sensitivity analysis across alternative Huber tuning constants to evaluate the stability of coefficient estimates in a Robust hurdle negative binomial model, used to model mobulid catch per unit effort for mobulid landings data collected in Threspuram Fishing Village and Tharuvaikulam Fishing Harbour, India in 2023. The red dotted lines represent the range of Huber constants (c) where the coefficients were most stable.

**Supplementary Table 2.2** Model estimates of the standard Hurdle Negative binomial model (HNB) for Catch per unit effort per fishing trip of mobulid bycatch in Tamil Nadu, India. For the count part, the positive catch per unit effort is the response variable and fishery characteristics (boat size, mesh size, fishing depth, fishing distance, and fishing days) as independent variables. For the zero part, boat size and mesh size are independent variables. Note: Significance indicated by asterisks p < 0.05 (*), p < 0.01 (**), p < 0.001 (***). This model was a poorer fit than the Robust Hurdle negative binomial model (RHNB), and hence was not chosen to explain catch per unit effort. We present the standard HNB output here to show the effects that existed before the outliers were down-weighted for maximum likelihood estimation in the RHNB model.

| **Count model coefficients (truncated negative binomial with log link)** | | | | |
|---|---|---|---|---|
| | **Estimate** | **Std. error** | **z value** | **p-value** |
| Intercept | -6.749 | 1.674 | -4.032 | < 0.001 *** |
| Boat size (Medium-scale) | 0.554 | 0.325 | 1.703 | 0.088 |
| Boat size (Large-scale) | 1.369 | 0.743 | 1.843 | 0.065 |
| Mesh size | 0.066 | 0.731 | 0.091 | 0.464 |
| Fishing depth | 0.116 | 0.462 | 0.252 | 0.644 |
| Fishing distance | 0.571 | 0.227 | 2.506 | <0.05 * |
| Soak hours | 0.333 | 0.261 | 1.279 | 0.200 |
| Fishing days | 0.696 | 0.168 | 4.126 | < 0.001*** |
| Log (theta) | -0.498 | 0.319 | -1.559 | 0.119 |
| **Zero hurdle model coefficients (binomial with logit link)** | | | | |
| | **Estimate** | **Std. error** | **z value** | **p-value** |
| Intercept | -4.290 | 0.292 | -14.647 | < 0.001 *** |
| Boat size (Medium-scale) | 0.282 | 0.171 | 1.644 | 0.100 |
| Boat size (Large-scale) | 1.007 | 0.487 | 2.068 | < 0.05 * |
| Mesh size | 0.081 | 0.032 | 2.520 | < 0.05 * |
| **Pearson residuals** | | | | |
| **Minimum** | **1st quantile** | **Medium** | **3rd quantile** | **Maximum** |
| -0.2893 | -0.2020 | -0.1756 | -0.1433 | 20.2488 |
